# Model-driven design allows growth of *Mycoplasma pneumoniae* on serum-free media

**DOI:** 10.1101/2019.12.12.873117

**Authors:** Erika Gaspari, Antoni Malachowski, Luis Garcia-Morales, Raul Burgos, Luis Serrano, Vitor A. P. Martins dos Santos, Maria Suarez-Diez

## Abstract

*Mycoplasma pneumoniae* is a slow-growing, human pathogen that causes atypical pneumonia. Because it lacks a cell wall, many antibiotics are ineffective, and vaccination is required. Due to its reduced genome and dearth of many biosynthetic pathways, this fastidious bacterium depends on rich, undefined medium for growth, which makes large-scale cultivation for vaccine production challenging and expensive.

To understand factors limiting growth, we developed a genome-scale, constraint-based model of *M. pneumoniae* called iEG158_mpn to describe the metabolic potential of this bacterium. We have put special emphasis on cell membrane formation to identify key lipid components to maximize bacterial growth. We have used this knowledge to predict and validate *in vitro* two serum-free media able to sustain growth.

Our findings also show that glycolysis and lipid metabolism are much less efficient under hypoxia; these findings suggest that factors other than metabolism and membrane formation alone affect the growth of *M. pneumoniae*.

Altogether, our modelling approach enabled us to optimize medium composition, capacitated growth in defined media and streamlined operational requirements, thereby providing the basis for stable, reproducible and less expensive vaccine production.

## Introduction

Mycoplasma are a small genus of bacteria belonging to the Mollicute class,^1^ comprising 124 species, 14 of which are human pathogens,^2^ while others infect farm animals, herd animals and pets.^3^ Most large-scale animal farms make use of antibiotics to fight and prevent infections, which often lead to development of antibiotics-resistant bacteria.^4–6^ Moreover, chronic infections caused by several Mycoplasmas are not immediately detectable. The use of effective vaccines against various Mycoplasmas in farm animals is essential to prevent wide spreading of these chronic infections^7^. It has been suggested that, despite being primarily considered a human pathogen, *Mycoplasma pneumoniae* (MPN) could potentially be used as a universal chassis to be deployed as single-or multi-vaccine in a range of animal hosts (www.mycosynvac.eu).

MPN causes atypical or walking pneumonia.^8^ With its highly reduced genome (816 kb), it is among the smallest self-replicating living organisms and has an exclusively parasitic lifecycle. This feature greatly determined its evolution towards minimal functions (e.g. absence of vitamins and lipids synthesis) and inability to detoxify metabolic waste (e.g. hydrogen peroxide), tasks for which this bacterium fully relies on the host.^9^ MPN primarily ferments glucose^10^ and uses most of its energy (71 to 88%) for non-growth associated tasks^11^. One of the key features of MPN during infections is its ability to evade the immune system: the unique characteristics of its membrane (in particular the high cholesterol contents) allows the mimicking of the host membrane.^12^ Moreover, the lack of a cell wall makes it resistant to antibiotics which target this structure.^13^

The development of vaccines based on MPN requires the large-scale production of this bacterium in minimal chemically defined medium. This is due to the high economic cost of rich media, the variable composition of animal-derived compounds^14^ and the possible presence of toxins, viruses or antigens that could reduce the sensitivity of immunological assays used on farms to screen for infection. ^15^ However, the production of MPN in defined media is a major challenge. For decades, the ability to culture MPN was reported to be possible only in serum-rich media^16^ due to a requirement for sterol components.^17^ Only in 2009 Yus et al. reported a defined medium based on the metabolic map reconstruction of MPN that allowed some growth of the bacterium but required daily changes of the medium^18,19^. This medium is not valid for large-scale growth needed during vaccine production. For these reasons, we focused our study on the model-driven design of a serum-free medium that could improve MPN growth rate.

We approached the study through metabolic modeling, a method widely used for genome-scale biochemical networks analysis.^20–27^. We previously described the development of a genome-scale, constraint-based model (GEM) of MPN metabolism, iJW145,^28^ and its use to carry out predictions related to growth rate and medium design, relying upon a biomass composition that had been formulated from literature,^3^ sequencing, proteomics and mass spectrometry data. Razin et al.^3^ reported the biomass of MPN is composed of 54-62% protein, 12-20% lipids, 3-8% carbohydrates, 8-17% RNA and 4-7% DNA. However, no novelty in the defined medium composition was reported^28^. Several studies show that lipids were the most important component limiting growth, mainly cholesterol, regulating membrane fluidity, and fatty acids.^1,29–32^ Determining the lipids composing the membrane of MPN is challenging, as it is strongly dependent on the composition of the growth medium, growth phase and culture conditions.^33^ Some key features remain invariant, such as the high cholesterol proportion, directly incorporated from the medium.^34^ The model iJW145 did not account in detail for lipids in its biomass composition^11,28^. Therefore, we integrated native lipid pathways of MPN, and reactions involved in the membrane formation to develop an updated GEM of MPN metabolism, called iEG158_mpn.

We aimed to describe the use of the energy available, taking into account not only cytosolic processes but also transmembrane transport. We used the model iEG158_mpn to design serum-free media for MPN and studied in detail how culture conditions affect the growth of MPN, focusing on chemical-physical aspects of the intracellular metabolism. Our study, in combination with the existing literature, suggests membrane formation and adaptability are key aspects to be investigated in relation with MPN growth, which appears to be limited by several factors.

## Materials and Methods

### Reconstruction of the genome-scale, constraint-based metabolic model

The metabolic model iJW145^28^ was extensively updated, manually curated and expanded to generate the genome-scale metabolic model iEG158_mpn. Reactions and metabolites have been annotated adding BiGG Models identifiers^35^ when possible. Reaction reversibility has been assessed through eQuilibrator,^36^ a thermodynamic calculator of the Gibbs energy released by a specific reaction at a set pH and ionic strength (in our case, respectively 7.0 and 0.1M). Reactions where changes in Gibbs energy comprised between −30 and 30 kJ/mol were considered reversible, otherwise irreversible. All model reactions have been manually verified using information from the MyMpn database.^37^ Information gathered from literature was used to modify the biomass synthesis reaction adding detailed information on lipid composition. Reactions for fatty acid integration into membrane lipid chains have been added. The genome-scale model iEG158_mpn is in SBML format and its syntax and consistency has been checked with SBML Validator.^38^ A list of all reactions and metabolites IDs is given in Supplementary Table S1. The model has been deposited in BioModels^39–41^ and assigned identifier MODEL1912060001. A python script to simulate growth with this model, imposing constraints and supplementing *in silico* the medium, is provided as Supplementary File S1.

### Model simulations and medium design

Pathways and whole-metabolism maps for fluxes visualization were constructed with the tool Escher,^42^ which graphically depicts a solution for maximum attainable growth rate, as computed by Flux Balance Analysis (FBA), with biomass yield, as a proxy for growth, as the objective function.^43^ FBA solves a linear programming problem, which is characterized by having more variables than equations, meaning more than one optimal solution can exist for the same objective function maximization. FBA computes a possible flux distribution leading to the optimal objective solution. The same optimal growth rate might be reached by several different reaction flux combinations. Flux Variability Analysis (FVA) explores the range of each metabolic flux at the optimal solution.^44^ Both FBA and FVA were run with biomass synthesis reaction as objective function to maximize, respectively, through the commands *optimize()* and *flux_variability_analysis()* of the Python tool *cobrapy*,^45^ version 0.5.11 (Python version 3.4.4). Exchange reactions in a genome-scale metabolic model allow metabolites to enter and egress the *in silico* network. Simulations with iEG158_mpn were performed by conserving the same flux constraints as have been experimentally determined: these assumed unlimited availability of the compounds which are not directly metabolized for energy or carbon and/or cannot be measured experimentally (vitamins, cofactors). Additional fatty acids that could be metabolized or integrated as membrane lipid chains (palmitic acid, palmitelaidic acid, stearic acid, oleic acid, linoleic and linolelaidic acid) have also been assumed to have unlimited availability as no related uptake rates were experimentally measured. Specifically, exchange reactions corresponding to metabolites whose uptake or secretion rate was not measured were set with bounds of ± 1000 when production and secretion was possible, with a lower bound of −1000 and an upper bound of 0 when only uptake from medium was possible and with a lower bound of 0 and an upper bound of 1000 when describing secretion.

Growth medium optimization consists of changing the bounds of exchange reactions to enable or limit uptake or secretion in order to find the combination of supplemented compounds that maximizes the flux through the biomass reaction, if this corresponds to maximizing growth rate. We tested the possible uptake of additional native compounds not previously considered in MPN growth media, but for which transporters are present in MPN genome, by adding the corresponding exchange reactions (Supplementary Table S1). To equilibrate the energetics in the model, we assigned a cost of 1 proton for the uptake of uncharged compounds, 1 ATP for the uptake of compounds with a charge of 1 or 2 and 2 ATPs for the one of compounds with charge greater than 2. To design a minimal chemically defined medium that optimizes MPN growth, we iterated through all the exchange reactions present to find the combination of bounds that lead to the highest biomass production: we have initially set all the exchange reactions’ lower bounds to −1000, allowing the metabolites of interest into the system. All the lower bounds of the exchange reactions were set to 0 one at a time, to be then repristinated to −1000 for the next iteration (next exchange reaction to close). When biomass yield was not changing in consequence, we assumed the component was not required for maximizing growth rate. Each time a component was removed from the list, a new set of iterations was started on all the other components. The minimal medium was obtained once all the components, when removed (i.e. when the correspondent exchange reaction’s lower bound was set to 0), were changing the biomass yield obtained by FBA. The relation between the growth rate and the doubling time has been computed using the exponential function e^µT^=2, where *µ* is the growth rate in divisions per unit of time and *T* is the doubling time.

To simulate hypoxia, we first ascertained, through Flux Balance Analysis, the minimal oxygen uptake rate required for optimal growth by iEG158_mpn. By allowing *in silico* an infinite supplementation of oxygen (“EX_O2_e” reaction lower bound = −1000), we ran FBA towards flux maximization of the biomass reaction. Constraining the growth rate to the maximal one, following FBA with as objective function “EX_O2_e” (minimization of oxygen uptake), we computed the minimal oxygen required. Once this value is established as the minimal oxygen uptake required for optimal growth, we reduced the oxygen supplementation (i.e. subsequent increments of “EX_O2_e” reaction lower bound from maximum oxygen uptake rate in baseline conditions towards 0) until reaching a biomass flux of 0, meaning no growth *in silico*. Following this approach, we extracted the minimum required oxygen to be supplemented *in silico* to allow growth. We then performed simulations of iEG158_mpn with an intermediate oxygen supplementation resulting from the rounded calculation of the mean of the minimal oxygen uptake required for optimal growth and the minimal oxygen required for allowing growth.

### *In silico* prediction of optimal medium lipids based on membrane composition

Optimal lipid composition of the medium was predicted according to the scheme in Figure 1. Each candidate medium lipid was integrated into the cytosol and subsequently into membrane compartment of biomass in three possible alternative ways: through direct integration, after being metabolized or after synthesis from precursors in the medium. Each lipid is considered carrying quantities of fatty acids, allowing the optimization not just of the membrane lipids but also of an experimentally determined fatty acids profile. Not only one but several proportions distributions of the membrane composition are considered, according to the ones detected in literature (as in our model: see Supplementary Table S2). Optimization of the lipids in the growth medium (see previous section) is then performed balancing both optimal fatty acid profile and variable membrane compositions, in order to predict a consensus medium composition able to allow growth under all the membrane configurations detected during laboratory cultivation.

**Figure 1.**
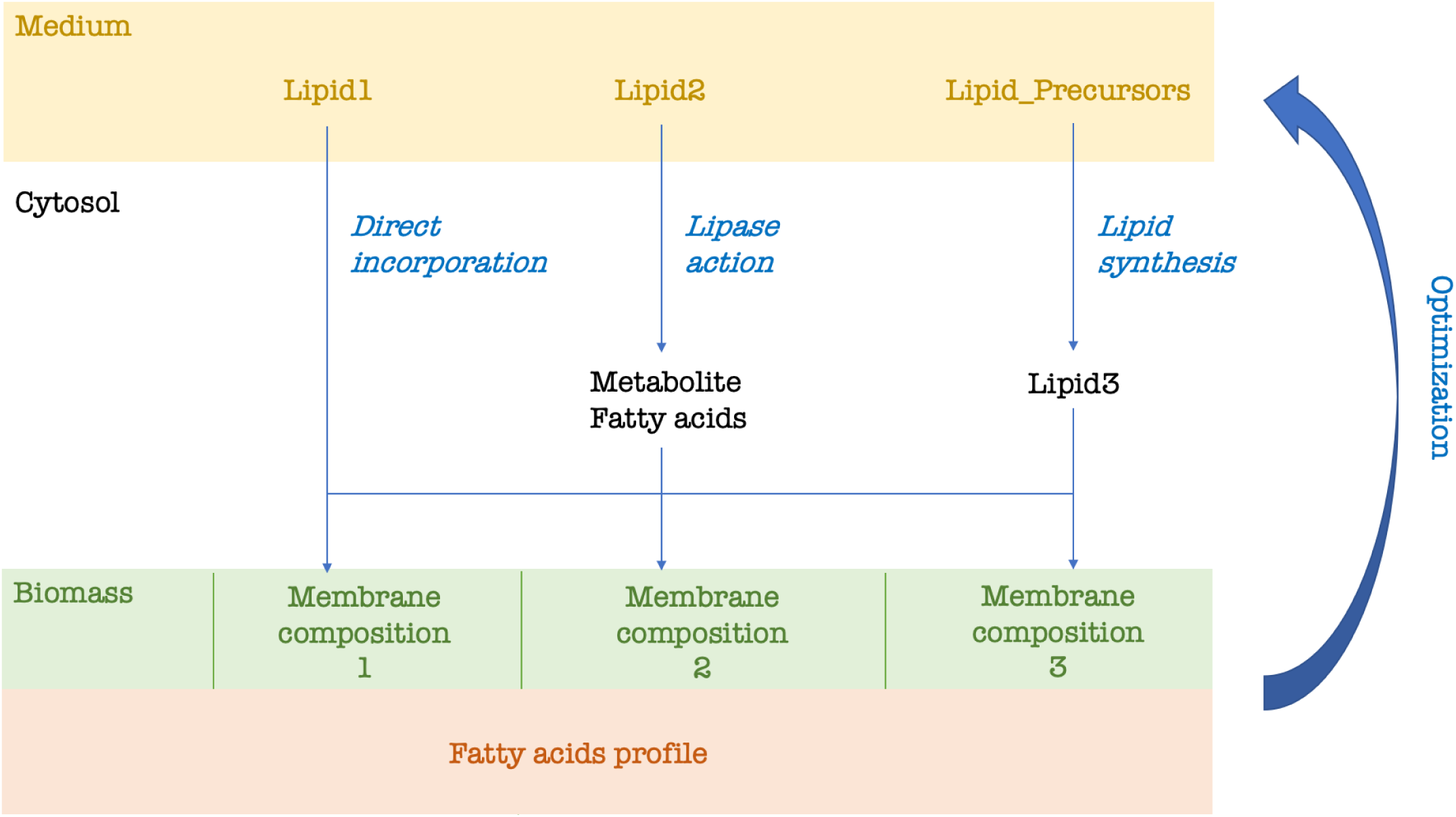
*In silico* optimization strategy for predicting optimal lipid composition of growth medium. All the possible lipids and precursors are provided in the medium for uptake. Each possible membrane composition that is found in the literature can be incorporated in the model as part of the biomass synthesis reaction. In our case, we considered three membrane compositions whose lipid proportions change according to the cholesterol percentage. Fatty acid profile can be found in literature (as in our model) or be experimentally determined. Lipid1 is known to be directly incorporated into the membrane, Lipid2 is metabolized in the cytosol and, whether a lipase acts, degraded into component head group and fatty acids. Lipid3 is synthetized in the cytosol from lipid precursors. Each lipid is integrated into the membrane according to the compositions previously reported in the literature. According to the optimal membrane composition and fatty acid profile, a consensus minimal lipid composition of the medium is predicted.

### Sequence alignment

By querying manually-curated sequences in the UNIPROT database,^46^ we verified the presence of lactate and acetic acid transporters and general monocarboxylic acid transporters in the MPN genome (Supplementary Files S2). PSI-BLAST algorithm^47^ was run with default parameters until convergence was reached to build a sequence profile of the transporters (Position Specific Scoring Matrix provided in Supplementary Files S2), then used to query within MPN genome for presence of these transporters. In addition, we searched for monocarboxylic transporters in MPN by running a BLASTp search with default parameters on all its transmembrane proteins, whose list was available on the *MyMpn* portal (August 2018),^37^ filtering for proteins containing transmembrane helices.

### Cultivation conditions for experimental validations of lipid requirements

*M. pneumoniae* strain M129 was grown in 75cm^2^ Tissue Culture Treated Flasks (Falcon) in a serum-free medium previously used for growing *Mesoplasma florum*.^48,49^ Palmitic acid, oleic acid and cholesterol were added to this medium at a final concentration of 16.5 µg mL^−1^, 20 µg mL^−1^ and 20 µg mL^−1^, respectively. Cells were grown and passaged three times in absence or presence of sphingomyelin and phosphatidylcholine at 20 µg ml^−1^ before performing any analysis. Each passage was performed by diluting the cells 50 times in fresh serum-free medium after two weeks of culture. Medium was replaced by fresh serum-free medium every 7 days.

Passage 3 cells DNA were measured by qPCR using oligonucleotides MPN628/5 (GCCATTTTGGATGGTTATGG) and MPN628/3 (GGTGACCCACTTCCGAGTTA) and the Luna® Universal qPCR Master Mix (New England Biolabs). The DNA quantification was performed using the LightCycler® 480 software (Roche). Cells were diluted in 15 mL to an equivalent of 0.6 pg mL^−1^ of DNA and distributed in aliquots of 0.2 mL in 96-well Tissue Culture Treated flat plates (Nunc). Every 12h, the medium from 6 wells were recovered in 1.5 mL tubes and centrifuged 15 min at 12 000g. 0.2 mL of nuclease-free water were added to the empty wells to lyse adherent cells by osmotic shock and then recovered and added to the previous pellet to lyse non-adherent cells. Finally, they were incubated 10 min at 98°C and stored at −20°C. After 120h, the DNA of each sample was measured by qPCR as previously described and the growth curve was then plotted and analysed using the R software (version 3.4.4).

*M. pneumoniae* strain M129 was grown in a 96-well plate format at 37°C in a serum-free medium developed by CRG. Growth curves comparing the performance of the serum-free medium containing phosphatidylcholine (PC), sphingomyelin (SMP) or both were recorded in a Tecan Spark plate reader by determining the growth index value, which is the ratio of absorbance at 430 nm and 560 nm of the culture medium.^19^ Total cell biomass obtained at the end of the growth curve (96h) was also determined by protein quantification using the Pierce BCA protein assay kit as previously described.^19^ Disclosure of the serum-free medium composition is prevented by third party agreements, but it may be available upon personal request, exclusively for validation purposes, under a material transfer agreement with CRG.

## Results

### Features of iEG158_mpn

The model iEG158_mpn of MPN metabolism has two compartments, extra-cellular and cytosolic, and consists of 490 reactions, 442 metabolites and 158 genes. Of the total reactions, 314 are gene-associated, 329 are cytosolic conversion reactions, 65 represent transport between the extracellular compartment and the cytosol and vice versa, and 96 are of exchange, meaning they allow uptake/production of compounds. Only 61 of the cytosolic reactions have a zero-flux range, meaning they are not active under the simulated conditions. Excluding the zero-flux reactions, 293 reactions have same minimum and maximum fluxes according to FVA, meaning in total 354 reactions with fixed flux. The high proportion of active reactions and the lack of flexibility in the usage of alternative reactions reflect the evolution of the organism in minimal genome. Minor cell components mainly consist of carbohydrates, amino acids and nucleotides. In the biomass equation, formulated according to Wodke et al.^11^, cofactors and vitamins, known to be essential for growth but with no available measurements, were symbolically included as traces in the biomass equation. Similarly, DNA repair and RNA turnover were considered in the biomass composition by slightly increasing their proportions. Protein turnover is modeled by the addition of degradation reactions,^50^ which is reflected in iEG158_mpn as the decomposition of a general protein compound into the essential amino acids and the associated ATP cost.

The very small size of MPN contributes to the increase in maintenance costs, as the high surface-to-volume ratio induces higher proton leakage through the membrane. The ATP maintenance reaction was determined in iJW145 by multiplying the number of ATPase complexes on the cell surface by the number of cells in a gram of biomass, so considering only the cytosolic side of the energy requirement. However, the reaction fluxes representation allowed the visualization of a flux of protons out of the cell, required for the cytosol de-acidification. Moreover, it is stated in Wodke et al.^28^ that the proportion of ATP used in cyclical lysis (which corresponds to total non-growth associated maintenance energy in iJW145) is about 76% at 24h after inoculation. We updated the non-growth-associated maintenance reaction to include both a cyclical ATP lysis component (33% of the total non-growth associated ATP consumption)(model reaction ID: “*IR08984*”) and a proton leakage component (66% of the total non-growth associated ATP consumption)(model reaction ID: “*protonLeak*”), as represented in Figure 2. Furthermore, this introduced a link between the maintenance equation and the proton production/consumption reactions.

**Figure 2.**
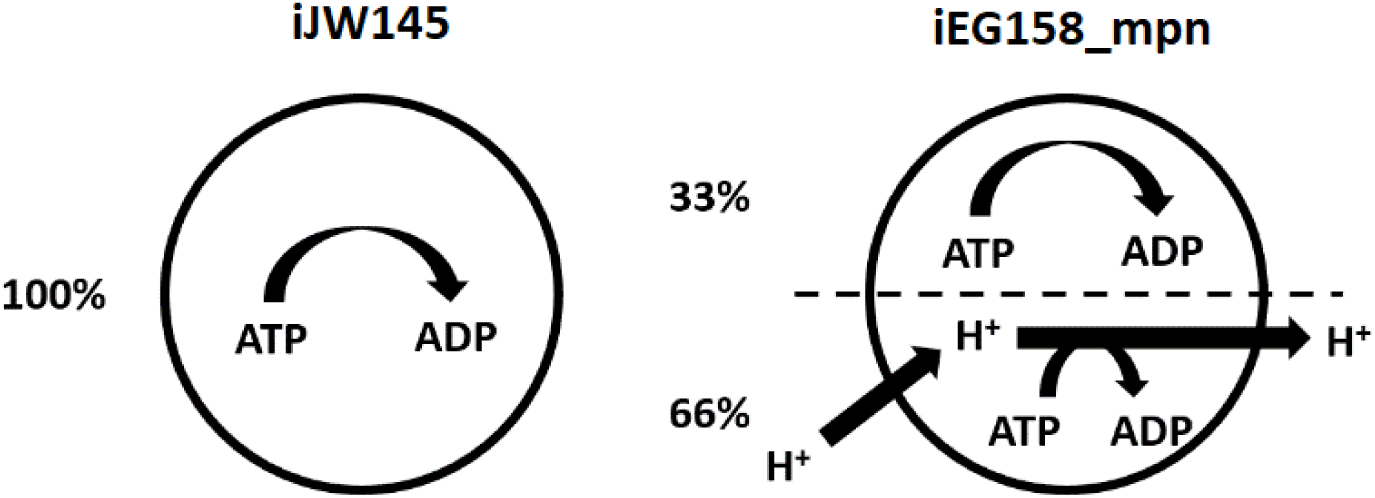
Representation of maintenance energy exploitation in GEM on *M. pneumoniae* left) iJW145 and right) iEG158_mpn. In iEG158_mpn simulations, ATP synthase is observed to run in reverse to allow efflux of protons for cytosol de-acidification. Maintenance reaction in iEG158_mpn was therefore updated to include a cyclical ATP lysis component (33%) and a proton leakage component (66%). The updated maintenance reaction in iEG158_mpn is compared to the one of iJW145, where only cyclical ATP lysis is assumed to be determinant.

The maintenance reaction in iEG158_mpn was further updated according to the results of the sequence alignment for monocarboxylic acids transporters discovery: no lactate or acetate proton symporter were identified to be present in MPN genome. The respective transport reactions (IDs “*LACLt”* and “*IR09888”*) were therefore updated in iEG158_mpn as irreversible, removing the proton symport and the maintenance costs adjusted to the maximum value allowing growth, computed as 10.46 mmol.g_DW_^−1^.h^−1^ of ATP (g_DW_=grams dry weight) used for non-growth-associated tasks at the quasi-steady state (24h time point).

ATP lysis and proton leakage are modelled in iEG158_mpn according to the described proportions of the maintenance energy usage: assuming the total maintenance cost, *M*_*t*_, equals to 10.46 mmol.g_DW_^−1^.h^−1^ of ATP, the energy fraction spent for proton leakage, *E*_*P*_, equals to 0.66 and the one used in ATP lysis, *E*_*L*_, equals to 0.33, the lower bounds *P*_*LB*_ and *L*_*LB*_ of the correspondent reactions (proton leakage and ATP lysis) are computed as [1] and [2]:

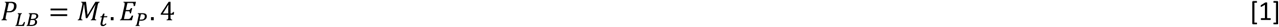

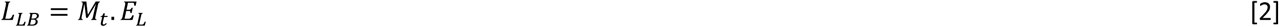

*P*_*LB*_ is multiplied by 4 as the reverse ATP synthase reaction removes four protons at a time.

We simulated the growth rate of MPN with iEG158_mpn supplementing *in silico* the medium predicted with model iJW145: the resulting growth rate is 0.053 divisions per hour at quasi-steady state of MPN (24h), correspondent to a doubling time of 13.4 hours, in line with the experimental evidences.

### Reconstruction of an optimal membrane lipids composition in iEG158_mpn

A model including only metabolic information has been shown not to be substantial to predict efficient strategies to maximize growth.^28^ Because membrane adaptability is one of the key features of MPN, we assumed that accounting for membrane formation in the model would give important insights into MPN growth. Focusing on membrane lipid components, we included in iEG158_mpn the pathways for lipids assembly^51^ and reconstructed the membrane lipid composition through an extensive literature search. Articles listed in this section were used to determine lipids percentages in the membrane and their acyl chains composition. Only lipids whose percentage in the membrane has been reported to be higher than 1% of the total lipid biomass were considered. Presence and proportions of the lipids are strongly dependent on the compound’s availability in the medium, therefore quantities must be interpreted as ranges.

One of the key features of the MPN’s membrane is the high proportion of cholesterol, from 35% to 50% of the total lipid fraction.^52^ *In vitro*, cholesterol is directly incorporated from the medium without being processed by the bacterium. For this reason, in iEG158_mpn, cholesterol has been included into the biomass composition without being metabolically processed. However, its proportion with respect to the other components in an optimal medium is easy to predict thanks to available literature, which indicates it constitutes about one third of the final membrane and about half of the lipid components of the membrane.

Most of the studies on MPN utilize Hayflick media, composed for 16% of horse serum, rich in sphingolipids and phosphatidylcholine.^53^ Sphingolipids-phosphatidylcholine ratio in MPN is 2.4,^54^ constituting on average 12% and 5% of the membrane lipids, respectively.^55^ However, sphingolipids have a good affinity for cholesterol, while phosphatidylcholine repels it, meaning their proportions in the MPN membrane are strongly correlated to the amount of cholesterol directly incorporated from the medium: a higher amount of cholesterol leads to the incorporation of a higher fraction of sphingolipids and a lower fraction of phosphatidylcholine^56^. The interrelation between these two lipids suggests their essentiality in the membrane is strongly related to their fatty acid chains, being the only lipids carrying the unsaturated fatty acid chain C18:2, whose presence might be determinant in MPN membrane. Moreover, sphingomyelin incorporated into the membrane has a lower proportion of fatty acid chains with 20 carbons or more in respect to the sphingolipids in the serum^57^, suggesting MPN preferentially incorporates in the membrane sphingomyelin with less than 20 carbons-fatty acid chains. MPN is a fatty acid and sterol auxotroph, but it can assemble phosphatidic acid, diacylglycerol, phosphatidylglycerol and glycolipids from provided fatty acids in the medium.^51^ Glycolipids constitute a range of 5-15% of the total membrane lipids,^58^ while the remaining percentage should be constituted by phosphatidylglycerol; phosphatidic acid and diacylglycerol, being lipid synthesis intermediaries, are not found in relevant proportions in the membrane.

The lipids, with their specific molecular weight, were included in the biomass equation to reflect the predicted membrane composition. We assumed a lipid composition is optimal when it confers optimal fluidity, and so permeability, to the membrane of MPN. Therefore, the acyl chains of the lipids constitute a critical aspect of our optimization.

With the aim of linking the membrane lipid profile to the availability of fatty acids in the medium, we reconstructed the acyl chains of the various lipids composing the membrane by considering that the sum of fatty acid chain fractions were previously determined by Wodke et al.^28^ and Worliczek et al.^57^; a comparison of the total fatty acid composition between our model and the ones available in literature is provided in Supplementary Figure S1. MPN tends to selectively incorporate palmitic acid over stearic acid in the membrane^32^, with a preference for saturated fatty acids, suggesting oleic and linoleic acids are incorporated in a lower proportion with respect to the amount found in the media. This indicates that the proportion of different acyl chains for de-novo assembled phosphatidic acids is conserved during assembly of downstream lipids phosphatidylglycerol, diacylglycerol and glycolipids. We assumed phosphatidylcholine could be degraded *in silico* into glycerol-3-phosphocholine and fatty acids, to represent the re-arrangements of the phospholipid’s acyl chains, which could be triggered by membrane lipases *in vivo* [REF]. Sphingomyelin, on the other hand, is in our model directly incorporated into the membrane without modifications: we therefore kept for this lipid the same fatty acid composition in the membrane as supplemented in the medium.

A representative average membrane lipids composition of MPN grown on serum rich medium is shown in Figure 3, with its respective and total fatty acid chains. In order to predict *in silico* the composition of serum-free medium that would maximize the growth of MPN, we added to the biomass equation the above representative medium composition, excluding those whose supplementation is linked to the addition of serum or would deviate from the optimal fatty acids proportions (e.g. cardiolipin). This way, simulating iEG158_mpn with as objective the maximization of the biomass flux (i.e. growth rate), we optimized for both growth and optimal membrane formation. We excluded phosphatidic acid from the biomass equation in iEG158_mpn and consider it solely as precursor of phosphatidylglycerol and glycolipids. Figure 4 consists of a visual representation of lipid pathways and membrane construction as they have been implemented in iEG158_mpn. The membrane is then composed of cholesterol, phosphatidylcholine, phosphatidylglycerol, glycolipids and sphingomyelin. We reconstructed their pathways within MPN metabolism and linked them to the medium components supplementing the related lipids. We included in the biomass equation the reported lipids, with their acyl chains, in their optimal proportion and maximize its flux to get the medium composition which enables growth.

**Figure 3.**
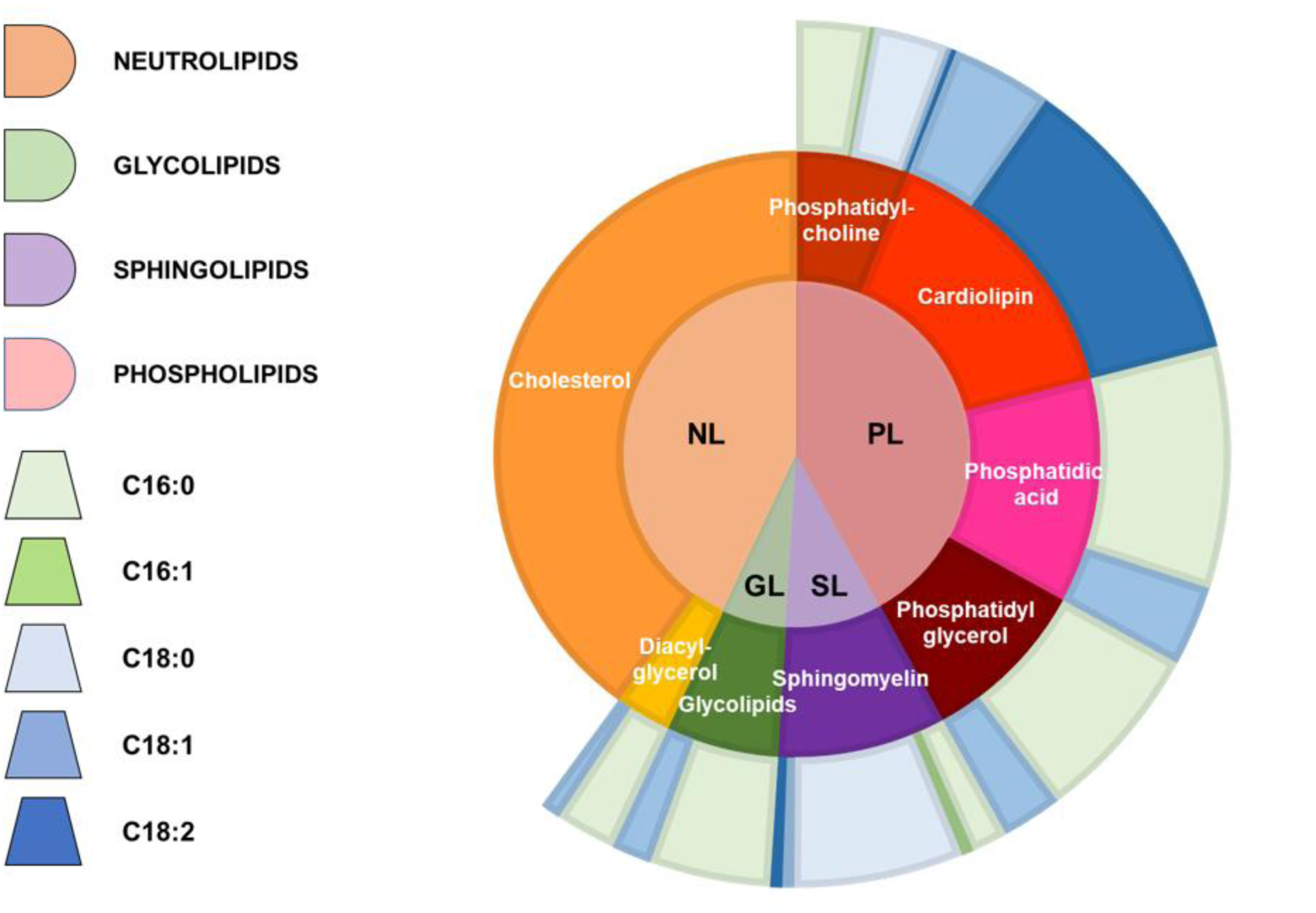
Reconstructed average composition of MPN membrane lipids and their respective fatty acid chains when the bacterium is grown on a serum-rich medium. The inner circle indicates lipid groups: neutrolipids (NL), glycolipids (GL), sphingolipids (SL) and phospholipids (PL). Lipids are represented in the intermediate circle as belonging to the different groups of the inner one; therefore, PL are constituted by phosphatidylcholine, cardiolipin, phosphatidic acid and phosphatidylglycerol, SL by sphingomyelin, GL by glycolipids and NL by cholesterol and diacyl-glycerol. Each lipid is then represented in the outer circle with their average fatty acid chains composition: the different proportions are made of palmitic acyl chains (C16:0), palmitoleic acyl chains (C16:1), stearic acyl chains (C18:0), oleic acyl chains (C18:1) and linoleic acyl chains (C18:2).

**Figure 4.**
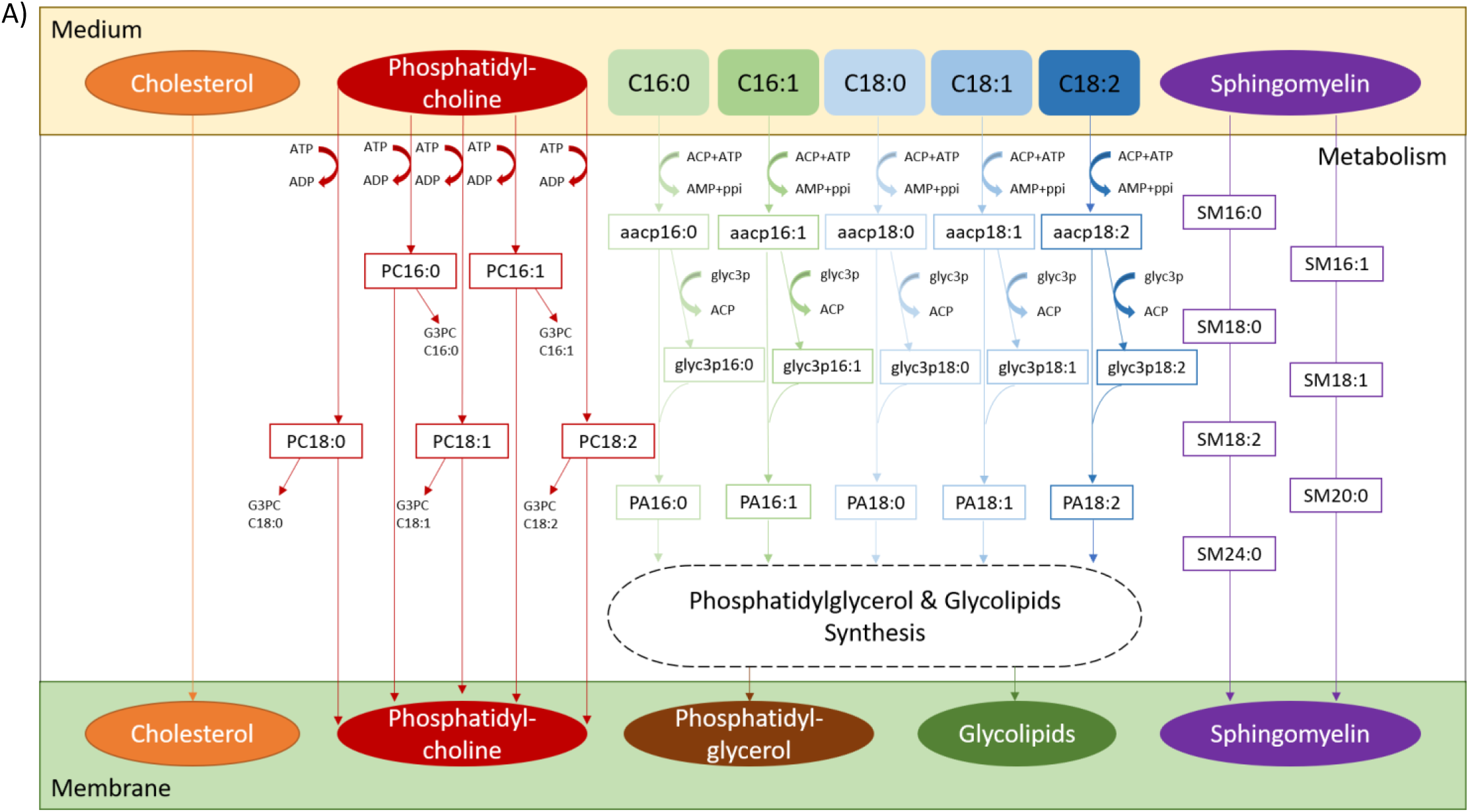
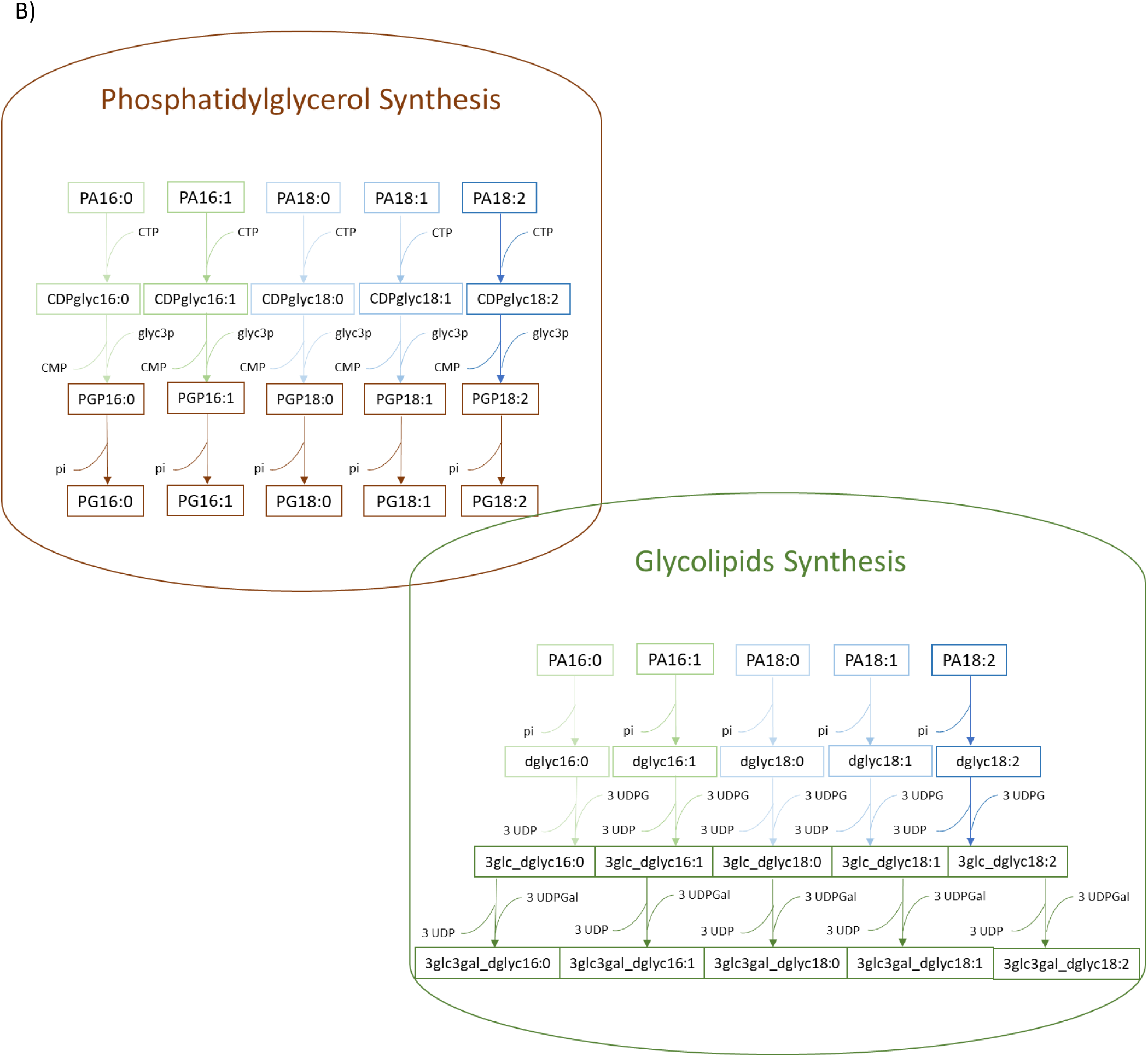
Implementation scheme of the lipid pathways and membrane formation in iEG158_mpn, when MPN is grown on serum-free medium. The wide variety of lipid species was simplified by considering all carry a representative acyl chain distribution instead of a mix with different acyl chain configurations. This simplification considerably reduces the complexity of the model keeping the same amount of quantitative information. A) Cholesterol is directly incorporated in the membrane as well as sphingomyelin (SM), for which all the different fatty acid chains proportions have been considered. Phosphatidylcholine is imported using ATP and either goes to build the membrane or it is degraded into its fatty acid chains and glycerol-phosphocholine (G3PC). All the fatty acids introduced in the medium (C16:0, C16:1, C18:0, C18:1, C18:2) will be linked to an acyl-carrier protein (ACP) at the expense of ATP. The ACP then releases the fatty acid chain to glycerol 3-phosphate (glyc3p) and the product, reacting with the ACP carrying fatty acids, leads to the production of phosphatidic acid (PA). B) PA then undergoes synthesis of phosphatidylglycerol (PG) or glycolipids.

The possible ranges of lipids and fatty acids proportions assuming growth in serum-free media are reported in Supplementary Table S2.

### Formulation of growth media incorporating specific lipids using a model-driven approach greatly increases MPN growth rate *in vitro*

We integrated into iEG158_mpn reactions involved in fatty acids, lipids and membrane formation pathways to reconstruct the membrane of MPN and predict essential fatty acids and lipids that, if added to the medium, could allow growth. As a result of this integration, the lipids in the biomass reaction of iEG158_mpn have been distributed as reported in Table 1.

**Table 1.** Lipid proportions in terms of percentage of total biomass and percentage of total biomass lipids.

| *LIPIDS* | PERCENTAGE OF TOTAL BIOMASS | PERCENTAGE OF TOTAL BIOMASS LIPIDS |
| --- | --- | --- |
| *Glycolipids* | 2.46 | 5.82 |
| *Cholesterol* | 18.13 | 42.92 |
| *Phosphatidylcholine* | 2.58 | 6.11 |
| *Phosphatidylglycerol* | 12.93 | 30.62 |
| *Sphingomyelin* | 6.14 | 14.53 |

From this membrane reconstruction, we have knowledge of the essential components that cannot be synthetize de-novo and therefore must be taken up from the medium. FBA shows the optimal components of the serum-free medium which maximize the growth of MPN, as shown in Table 2.

**Table 2.** Predicted media components to maximize MPN growth rate.

| CARBON SOURCES | VITAMINS | NUCLEOTIDES | AMINO ACIDS |
| --- | --- | --- | --- |
| Glucose | Nicotinic acid | Adenine | Alanine |
| Glycerol | Thiamin | Guanine | Arginine |
|  | Pyridoxal | Uracil | Asparagine |
|  | Riboflavin | Thymine | Cysteine |
|  | Choline | Cytidine | Glutamine |
|  | Folic acid | Deoxycytidine | Glycine |
|  | Pantetheine |  | Histidine |
|  | Adenosine 3',5'-bisphosphate |  | Isoleucine |
|  |  |  | Leucine |
|  |  |  | Lysine |
|  |  |  | Methionine |
|  |  |  | Phenylalanine |
| **LIPIDS** |  |  | Proline |
| Cholesterol |  |  | Serine |
| Palmitic acid |  |  | Threonine |
| Oleic acid |  |  | Tryptophan |
| Sphingomyelin |  |  | Tyrosine |
| Phosphatidylcholine |  |  | Valine |

The result about the components directly impacting the metabolism is consistent with the one previously listed in a serum-free defined medium that allowed weak growth^19^ and adds lipids components (sphingomyelin and phosphatidylcholine), essential for the membrane formation and stabilization. This led to the design of two experiments for *in vitro* assessment; results shown in Figure 5 and Figure 6 prove the determinant contribution of the synergy between sphingomyelin and phosphatidylcholine for MPN growth *in vitro*.

**Figure 5.**
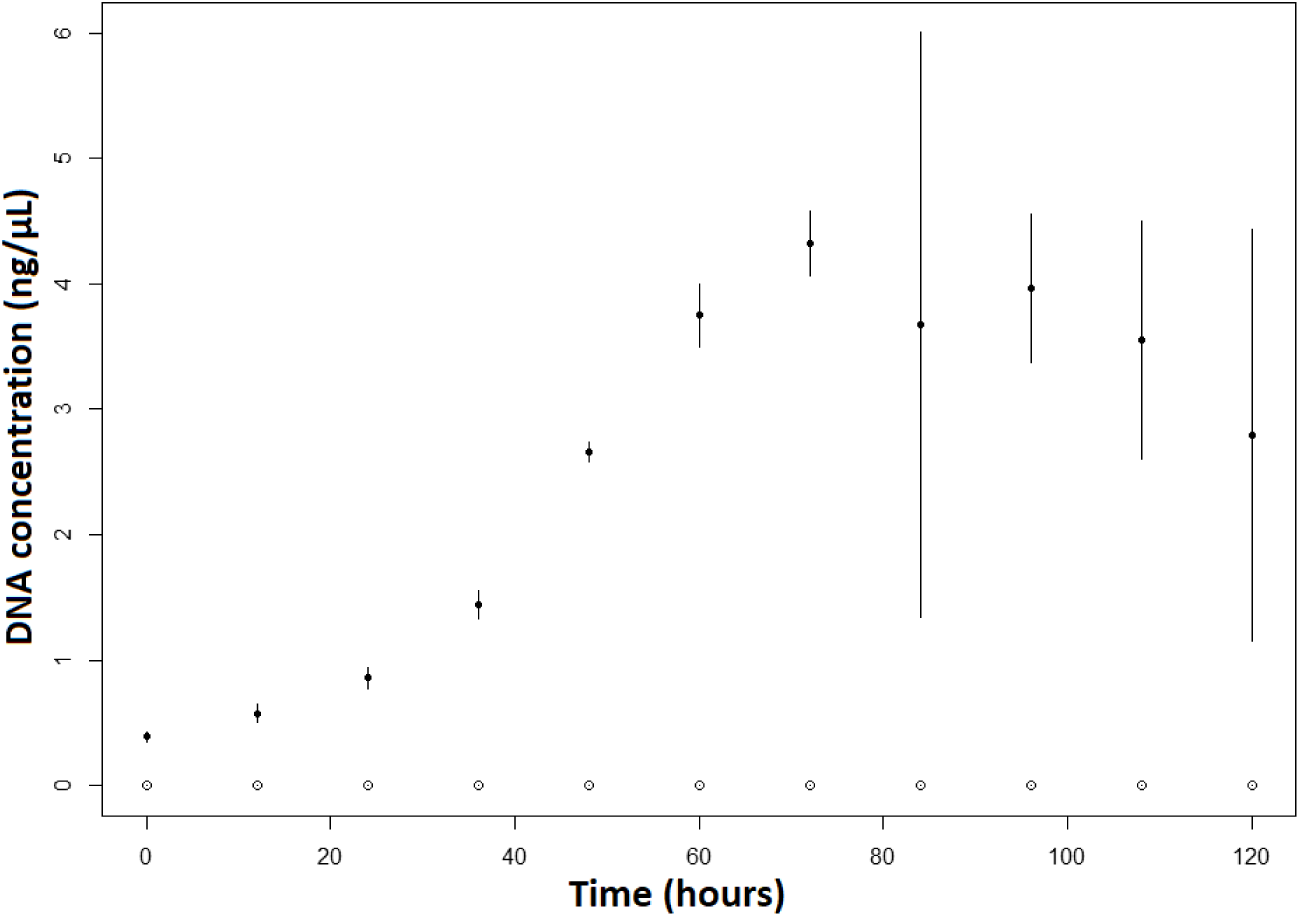
Growth of MPN cells in the serum-free *Mesoplasma florum* medium measured by genomic DNA quantification by qPCR. Full dots show the growth of MPN in presence of sphingomyelin and phosphatidylcholine. Empty dots show the growth of MPN in absence of both lipids. Bars represent standard deviation. Measurements represent the average of 6 replicates.

**Figure 6.**
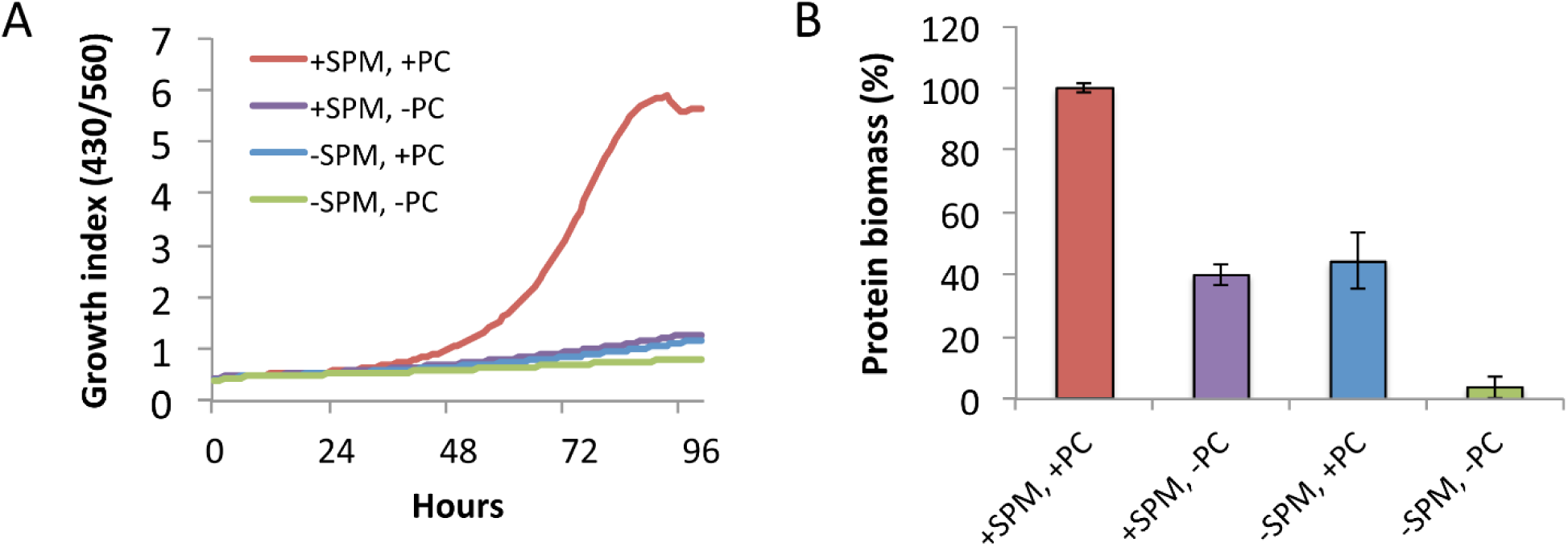
Impact of phosphatidylcholine and sphingomyelin on *M. pneumoniae* cell growth using the serum-free CRG medium. (A) Growth curve analysis determined by the 430/560 absorbance rate index comparing cell growth after adding phosphatidylcholine (PC) and sphingomyelin (SMP) individually or in combination in a medium free of serum. (+/-) indicates presence or absence of the indicated phospholipid. (B) Protein biomass measurement at 96 h, corresponding to the end of the growth curve shown in panel A. Data represent the mean ± standard deviation of two replicates.

Carbon sources, vitamins, nucleosides and amino acids components computed by our analysis are the same as determined by previous studies^18^. Our analysis pin-points, in terms of percentages of total lipid mass in the membrane, the three key lipid components (cholesterol, sphingomyelin and phosphatidylcholine) that were integrated in iEG158_mpn as follows: the most enriched components must be cholesterol, constituting 35-50% of the total lipids; sphingomyelin and phosphatidylcholine proportions are adjusted accordingly, as sphingomyelin has high affinity for cholesterol, while phosphatidylcholine has a low one.^56^ Therefore, they constitute respectively 9-15% and 6-10% of the total lipids. Figure 7 shows the interrelation between cholesterol, phosphatidylcholine and sphingomyelin (or in general sphingolipids) proportions in MPN membrane as simulated by iEG158_mpn. Sphingomyelin is preferably incorporated carrying an 18-carbon acyl chain and, therefore, its essentiality might be linked to it being the only one carrying the acyl chain C18:2.

**Figure 7.**
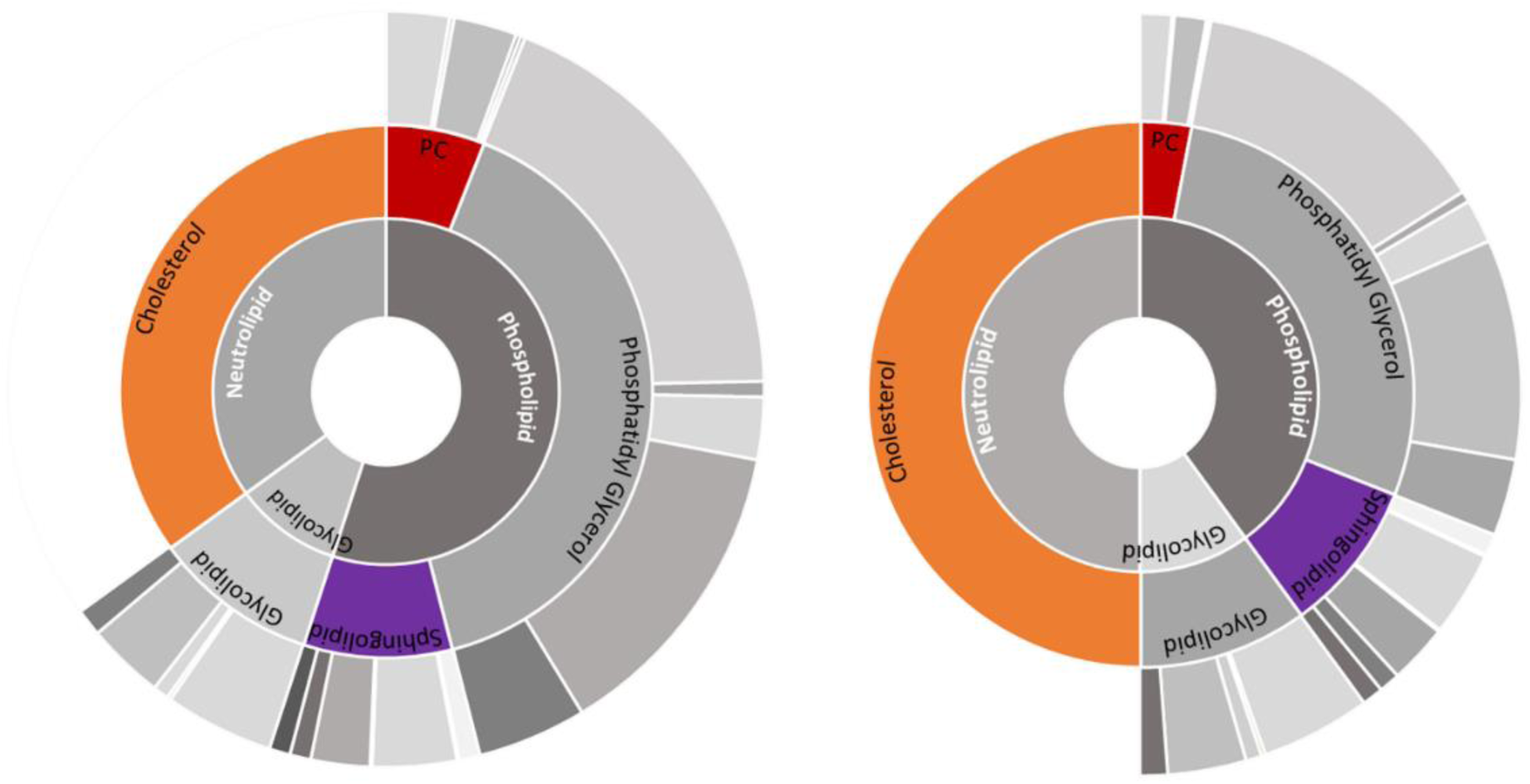
Interrelation between cholesterol, sphingomyelin and phosphatidylcholine proportions in the membrane of MPN as simulated by iEG158_mpn when grown *in silico* on the predicted optimal serum-free medium. A) If cholesterol constitutes 35% of the total membrane lipids, phosphatidylcholine (PC) proportion is up to 10%, while sphingomyelin/sphingolipids is reduced at its minimum to 12%. B) If cholesterol constitutes 50% of the membrane lipids, phosphatidylcholine (PC) proportion decreases to 6%, while sphingomyelin increases to 15%. Outer cycle represents the average proportions of acyl chains carried by each lipid, as also showed in Figure 3: the grey colors represent, in order of shading from the lightest to the darkest, C16:0, C16:1, C18:0, C18:1, C18:2.

The essential fatty acids to be incorporated in a serum-free medium are palmitic (C16:0) and oleic (C18:1) acids, as they are the preferred components of phosphatidic acid in MPN and its downstream products phosphatidylglycerol and glycolipids. Stearic acid (C18:0) presence is important depending on phosphatidylcholine proportion: once phosphatidylcholine is incorporated, its unsaturated fatty acid chains are replaced with saturated ones, resulting in the membrane di-saturated phosphatidylcholine.^59^ Moreover, supplementation of phosphatidylcholine might be important not only as membrane component but also as source of those fatty acids whose presence has been detected in lower percentage with respect to palmitic and oleic acids: linoleic acid (C18:2) and palmitoleic acid (C16:1).

### Effect of hypoxia on MPN metabolism

iEG158_mpn uses BiGG Models identifiers, so Escher can be applied to visualize change of fluxes when simulating different environmental conditions: Figure 8 shows flux changes for each reaction in part of the glycolysis pathway of the metabolic network when oxygen availability is reduced from minimal required uptake rate for maximal growth (7.54 mmol.g_DW_^−1^.h^−1^) to 6 mmol.g_DW_^−1^.h^−1^, as no growth occurs in complete absence of oxygen.60 Our model predicts absence of growth for oxygen uptake rates lower than 4.81 mmol.g_DW_^−1^.h^−1^. With no oxygen limitation and glucose uptake rate of 5.11 mmol.g_DW_^−1^.h^−1^, the model predicts MPN uptakes 7.54 mmol.g_DW_^−1^.h^−1^ of oxygen. The growth rate under hypoxic stress, in respect to baseline conditions, is limited by about 2.3-fold. FVA results for all the reaction fluxes in baseline and hypoxia conditions are reported in Supplementary Table S3, where the reactions included in Figure 8 are highlighted. Here the fluxes are represented in terms of differences between hypoxia and baseline conditions, given that the solutions for the flux distributions are unique. Despite the overall fluxes’ distribution being slightly reduced under oxygen limitation, the number of zero-flux reactions in this condition is 84. Excluding these, 299 reactions have same minimum and maximum allowed fluxes. The hyper-reduced size of the FVA solution space on both baseline and hypoxia, and so their unique solutions of the related FBA matrix, allow the representation of flux differences between the two conditions.

**Figure 8.**
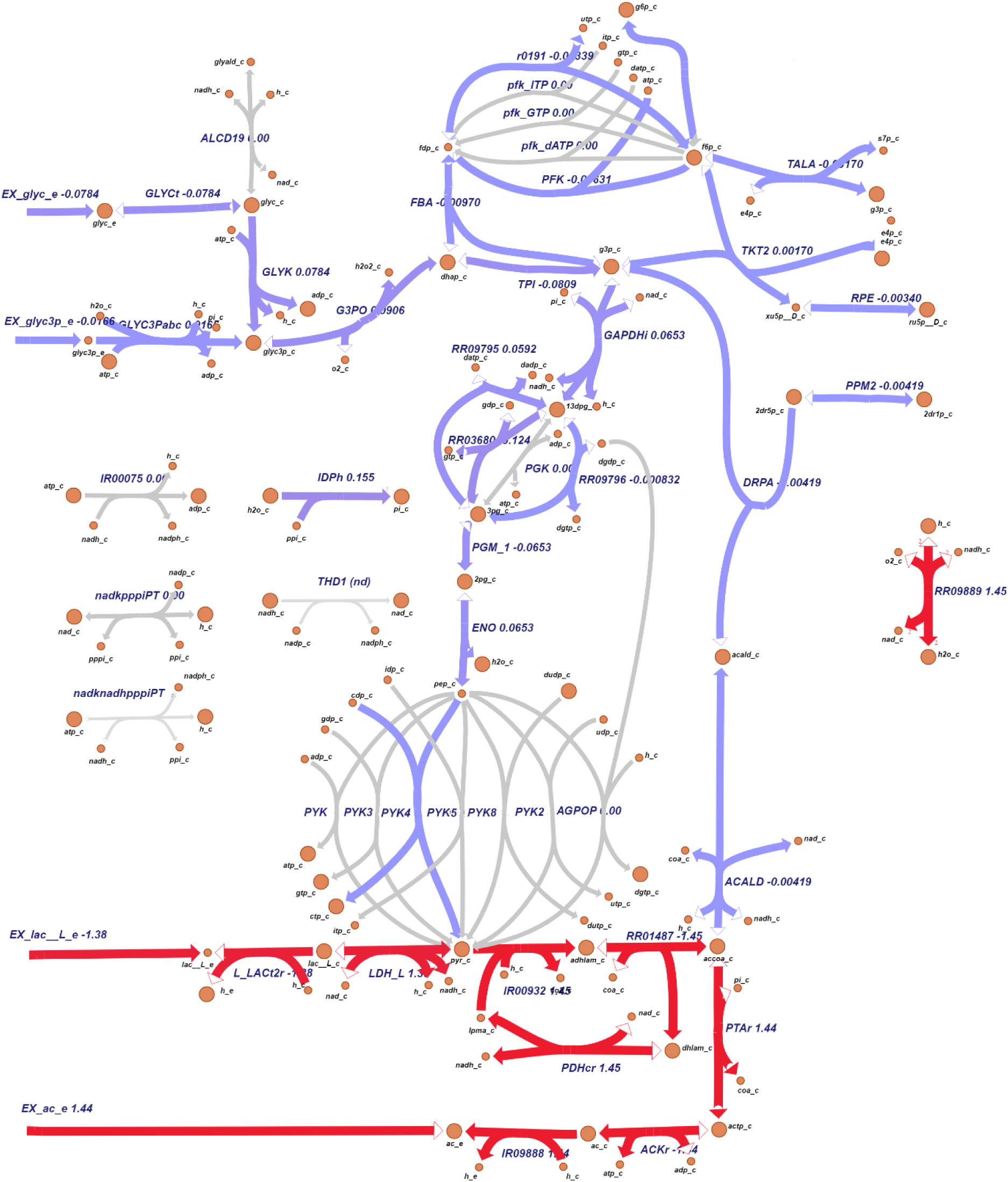
Visualization of flux differences in glycolysis reaction when MPN is under hypoxia respect to baseline oxygen availability (in mmol.g_Dw_^−1^.h^−1^). Purple is indicative of a small difference (absolute value of flux change < ±1) and red of a more considerable one (absolute value of flux change ≥ ±1). Grey arrows represent reactions that are unused or whose flux does not change when oxygen availability is reduced to 6 mmol.g_DW_^−1^.h^−1^.

Combining the information from FVA and the fluxes visualization, some conclusions on MPN under hypoxia can be drawn. Not considering the zero-flux reactions, only 18 (Supplementary Table S3) show the same range of fluxes comparing the two situations (baseline and reduced oxygen) in iEG158_mpn, assuming proton leakage and maintenance energy (ID: “IR08984”) do not change. All the lipid synthesis reactions show a consistent flux reduction, as well as the reactions related to the carbon degradation pathways and the NAD kinase. Imports of some amino acids and nucleobases (i.e. thymine, ID: “IR09855”) show a flux decrease when oxygen is reduced. Contrarily to the general tendency, deoxyuridine kinase and phosphorylase show a higher flux respect to the baseline situation. CO_2_, H_2_O_2_, acetic acid and lactic acid productions increase under hypoxia.

The whole-metabolism visualization of reaction flows under baseline conditions and when oxygen availability is reduced are reported in Supplementary Files S3.

## Discussion

Owing to its fastidious requirements, *Mycoplasma pneumoniae* has been thus far reported to grow on media containing serum^61,62^ and only poorly in defined medium^18^, furthermore requiring daily changes. Hereby, by using *in silico* modeling approaches based on literature data, we propose two media formulations that allow MPN to grow at a non-negligible rate, and we suggest that the poor growth reported in the previously defined medium was especially due to the absence of sphingomyelin and phosphatidylcholine. This prediction was validated by removing these phospholipids from the two different serum-free media formulations. The absence of animal-derived elements in these media facilitates the large-scale production of this microorganism for medical and pharmaceutical purposes, thanks to the reduced costs, greater reproducibility, definite composition and lack of unexpected serum-plate-agglutination (SPA) antigens^15,63^ and other unpredictable contaminants. Our study shows that supplementation of sphingomyelin and phosphatidylcholine to two serum-free media, together with cholesterol, enables growth of MPN. We also show that the different incorporation of these lipids depends on their availability in the medium. The MPN membrane is built on available resources and it is strongly variable according to the environmental conditions. The lipid composition of the membrane affects its permeability^64^ and curvature^65^. For instance, a sphingomyelin-cholesterol packing results in a very tight membrane^66^. Therefore, the key medium lipids herein found lead to the hypothesis that one of the main limitations of MPN growth resides in its membrane composition, fluidity and permeability. This is, in turn, determined by the proportion of acyl chains in the membrane lipids. While the requirement of cholesterol for growth has been well known for decades,^30,67^ phosphatidylcholine supplementation in a medium for MPN has been first reported in 2009^19^. Sphingomyelin, in contrast and to the best our knowledge has not been specifically reported as to be an essential component in media deployed to grow *Mycoplasma pneumoniae*.

Having established that lipids supplementation is a key aspect for MPN growth, the way they are supplemented might be critical: MPN cultures where lipids are supplemented as cholesterol-phosphatidylcholine vesicles show a similar growth rate of the ones in serum-rich media.^68^

Medium composition is not the only growth-limiting factor of MPN growth *in vitro*: batch cultures may result in limitation in oxygen supply as a result of gradients that arise due to biofilm formation. Our analysis shows oxygen level is critical, affecting the whole metabolism, with only a few reaction fluxes that remain intact under hypoxic conditions in respect to baseline conditions. Low oxygen not only pushes the lactate dehydrogenase towards the production of lactate, but also critically affects the whole glycolytic pathway, with a reduction of sugar uptake and ATP production. More NADH is consumed and less is regenerated by the GADPH enzyme, which represents one of the main bottlenecks of the metabolic network. The production and accumulation of compounds such as lactate and acetate augments, enhancing toxicity and acidifying the medium when exported: it is observed that the doubling time of MPN in batch culture increases over time due to the decrease in media pH^28^ and the accumulation of toxic waste compounds, namely acetic and lactic acid. Cells growing in an acidified medium have a different membrane composition,^33^ suggesting that the maintenance of an optimal media pH is also critical for optimal membrane lipids composition. It is, however, still unclear whether the adaptability of MPN membrane resides in its genome-coding function, if it is a chemical-physical response of the lipid bilayer to pH changes or if it is a combination of these two aspects.

The cytosol of Mycoplasmas does not maintain a constant pH relative to the outside environment, with the risk of being unable to maintain the proton gradient required for trans-membrane transport^69^ and to alter the internal biochemical state. Our model iEG158_mpn improves iJW145 by not only integrating the membrane, the specific lipids with their pathways and carried acyl chains, but also rearranging the ATP distribution in the maintenance energy, taking into account the effort made by MPN while de-acidifying the cytosol: a cytosolic pH decrease is equivalent to an increase in the H_3_O^+^ intracellular concentration, which might alter reactions involving H^+^ as reactant, for instance ATP hydrolysis.^70^ This might be a cause of high energy maintenance costs and reduced growth rate. For this reason, a study on the effects of the intracellular pH change on MPN metabolism would be of main interest.

A study of MPN under different environmental conditions would not only provide insights into the way it reacts to pH changes, but also disclose the strategies bacteria with a reduced genome use to exploit the host’s resources to accomplish essential tasks, whose capabilities it has lost during the course of evolution: our analysis highlights a lack of metabolic versatility, reflected by an *in silico* deficiency of flux variability, possibly due to an extremely reduced metabolic network size.

Altogether, our integrated approach, demonstrates the usefulness of modeling resources for exploring the metabolic capacity of mycoplasmas, and the use of these insights to rationally influence, modulate and design conditions for optimized performance.

## Acknowledgements

This work has received funding from European Research Council (ERC) under the European Union’s Horizon 2020 research and innovation program under grant agreement n. 634942 (MycoSynVac) and n. 670216 (MYCOCHASSIS).

## Author Contributions

EG, VAPMdS and MS-D conceived the project. EG and AM developed the model with suggestions from MS-D and LS. EG and AM performed modeling, simulations, calculations and analysis. LG-M performed serum-free *Mesoplasma forum* medium experiments. RB performed serum-free CRG medium experiments. EG wrote the manuscript with input of MS-D, AM, VAPMdS, LS, LG-M and RB. VAPMdS and MS-D supervised the project. All authors read and approved the manuscript.

## References

1. Razin, S. The mycoplasmas. Microbiol. Rev. 42, 414–470 (1978).

2. Hayflick, L., & Chanock, R.M.. Mycoplasma species of man. Bacteriol. Rev. 29, 185–220 (1965).

3. Razin, S., Yogev, D. & Naot, Y. Molecular biology and pathogenicity of mycoplasmas. Microbiol. Mol. Biol. Rev. 62, 1094–156 (1998).

4. Tilman, D., Cassman, K. G., Matson, P. A., Naylor, R. & Polasky, S. Agricultural sustainability and intensive production practices. Nature 418, 671 (2002).

5. Cabello, F. C. Heavy use of prophylactic antibiotics in aquaculture: a growing problem for human and animal health and for the environment. Environ. Microbiol. 8, 1137–1144 (2006)

6. Van Boeckel, T. P. et al. Global trends in antimicrobial use in food animals. Proc. Natl. Acad. Sci. 112, 5649–5654 (2015).

7. Nicholas, R. A. J., Ayling, R. D. & Stipkovits, L. P. An experimental vaccine for calf pneumonia caused by Mycoplasma bovis: clinical, cultural, serological and pathological findings. Vaccine 20, 3569–3575 (2002).

8. Grayston, J. T. et al. Mycoplasma pneumoniae Infections: Clinical and Epidemiologic Studies. JAMA 191, 369–374 (1965).

9. Collier, A. M. & Clyde, W. A. Relationships Between Mycoplasma pneumoniae and Human Respiratory Epithelium. Infect. Immun. 3, 694 LP–701 (1971).

10. Miles, R. J., Taylor, R. R. & Varsani, H. Oxygen uptake and H202 production by fermentative Mycoplasma spp. J. Med. Microbiol., 34, 219–223 (1991).

11. Wodke, J. A. H. et al. Dissecting the energy metabolism in Mycoplasma pneumoniae through genome-scale metabolic modeling. Mol. Syst. Biol. 9, (2013).

12. Razin, S. & Jacobs, E. Mycoplasma adhesion. J Gen Microbiol 138(3), 407–422 (1992).

13. Waites, K. B. & Talkington, D. F. Mycoplasma pneumoniae and Its Role as a Human Pathogen. Clinical Microbiology Reviews, 17, 697–728 (2004).

14. Merten, O. W. Safety issues of animal products used in serum-free media. Dev. Biol. Stand. 99, 167—180 (1999).

15. Ahmad, I., Kleven, A. S. H., Avakian, A. P. & Glisson, J. R. Sensitivity and Specificity of Mycoplasma gallisepticum Agglutination Antigens Prepared from Medium with Artificial Liposomes Substituting for Seru. American Ass. 32, 519–526 (2018).

16. Laidlaw, P.P. & Elford, F.R.S. A New Group of Filterable Organisms. Royal Society, 120(188), (1936).

17. Edward, B. Y. D. G. F. F. & Fitzgerald, W. A. The Isolation of Organisms of the Pleuropneumonia Group from Dogs. J. Gen. Microbiol. 5, 566–575 (1951).

18. Yus, E. et al. Impact of Genome Reduction and Its Regulation. Science (80-.). 326, 1263 (2010).

19. Yus, E. et al. Impact of genome reduction on bacterial metabolism and its regulation. Science (80-.). 326, 1263–1268 (2009).

20. Price, N. D., Papin, J. A., Schilling, C. H. & Palsson, B. O. Genome-scale microbial in silico models : the constraints-based approach. TRENDS in Biotec. 21, 162–169 (2003).

21. Westerhoff, H. V & Palsson, B. O. The evolution of molecular biology into systems biology. Nature Biotec. 22, 1249–1252 (2004).

22. Edwards, J. S., Covert, M. & Palsson, B. Metabolic modelling of microbes: the flux-balance approach. Environ. Microbiol. 4, 133–140 (2002).

23. Mardinoglu, A. et al. Genome-scale metabolic modelling of hepatocytes reveals serine deficiency in patients with non-alcoholic fatty liver disease. Nat. Commun. 5, 3083 (2014).

24. Batstone, D. J., Hülsen, T. & Oehmen, A. Metabolic modelling of mixed culture anaerobic microbial processes. Curr. Opin. Biotechnol. 57, 137–144 (2019).

25. Shene, C., Asenjo, J. A. & Chisti, Y. Metabolic modelling and simulation of the light and dark metabolism of Chlamydomonas reinhardtii. Plant J. 96, 1076–1088 (2018).

26. Razmilic, V., Castro, J. F., Marchant, F., Asenjo, J. A. & Andrews, B. Metabolic modelling and flux analysis of microorganisms from the Atacama Desert used in biotechnological processes. Antonie Van Leeuwenhoek 111, 1479–1491 (2018).

27. Pfau, T. et al. The intertwined metabolism during symbiotic nitrogen fixation elucidated by metabolic modelling. Sci. Rep. 8, 12504 (2018).

28. Wodke, J. A. H. et al. Dissecting the energy metabolism in Mycoplasma pneumoniae through genome-scale metabolic modeling. Mol. Syst. Biol. 9, 653–653 (2014).

29. Kurzepa, H., Flinton, L. & Vandemark, P. J. Growth of Parasitic Mycoplasma Without Serum or Serum Fraction. J. Bacteriol. 99, 908–909 (1969).

30. Razin, S. & Tully, J. G. Cholesterol Requirement of Mycoplasmas. J. Bacteriol. 102, 306–310 (1970).

31. Rottem, S. & Kahane, I. Mycoplasma cell membranes. Springer Science & Business Media 20, (2012).

32. Leon, O. & Panos, C. Long-chain fatty acid perturbations in Mycoplasma pneumoniae. J. Bacteriol. 146, 1124 LP–1134 (1981).

33. Pollack, J. D., Somerson, N. L. & Senterfit, L. B. Effect of pH on the immunogenicity of Mycoplasma pneumoniae. J. Bacteriol. 97, 612–619 (1969).

34. Dahl, J. S. & Dahl, C. E. Effect of Cholesterol on Macromolecular Synthesis and Fatty Acid Uptake by Mycoplasma capricolum. J. Biological Chemistry. 256, 87–91 (1981).

35. King, Z. A. et al. BiGG Models: A platform for integrating, standardizing and sharing genome-scale models. Nucleic Acids Res. 44, D515–D522 (2015).

36. Flamholz, A., Noor, E., Bar-even, A. & Milo, R. eQuilibrator — the biochemical thermodynamics calculator. Nucl. Acids Res. 40, 770–775 (2012).

37. Cozzuto, L. et al. MyMpn : a database for the systems biology model organism Mycoplasma pneumoniae. Nucl. Acids Res. 43, 618–623 (2015).

38. Hucka, M. et al. The systems biology markup language (SBML): a medium for representation and exchange of biochemical network models. Bioinformatics. 19, 524–531 (2003).

39. Le Novère, N. et al. BioModels Database: a free, centralized database of curated, published, quantitative kinetic models of biochemical and cellular systems. Nucleic Acids Res. 34, D689–D691 (2006).

40. Li, C. et al. BioModels Database: An enhanced, curated and annotated resource for published quantitative kinetic models. BMC Syst. Biol. 4, 92 (2010).

41. Chelliah, V. et al. BioModels: ten-year anniversary. Nucleic Acids Res. 43, D542–D548 (2014).

42. King, Z. A. et al. Escher : A Web Application for Building, Sharing, and Embedding Data-Rich Visualizations of Biological Pathways. PLOS Comp. Bio. 1–13 (2015).

43. Orth, J. D., Thiele, I. & Palsson, B. Ø. What is flux balance analysis? Nat. Publ. Gr. 28, 245–248 (2010).

44. Gudmundsson, S. & Thiele, I. Computationally efficient flux variability analysis. BMC Bioinformatics 11, 489 (2010).

45. Ebrahim, A., Lerman, J. A., Palsson, B. O. & Hyduke, D. R. COBRApy: COnstraints-Based Reconstruction and Analysis for Python. BMC Syst. Biol. 7, 74 (2013).

46. Magrane, M. & Uniprot Consortium. UniProt Knowledgebase : a hub of integrated protein data. Database. 2011, 1–13 (2011).

47. Altschul, S. F. et al. Gapped BLAST and PSI-BLAST : a new generation of protein database search programs. Nucl. Ac. Res. 25, 3389–3402 (1997).

48. Mccoy, R. E. et al. Acholeplasma florum, a New Species Isolated from Plants. Int. J. Syst. Evol. Microbiol. 34, 11–15 (1984).

49. Tully, J. G., Bové, J. M., Laigret, F. & Whitcomb, R. F. Notes: Revised Taxonomy of the Class Mollicutes: Proposed Elevation of a Monophyletic Cluster of Arthropod-Associated Mollicutes to Ordinal Rank (Entomoplasmatales ord. nov.), with Provision for Familial Rank To Separate Species with Nonhelical Morphology (Entomoplasmataceae fam. nov.) from Helical Species (Spiroplasmataceae), and Emended Descriptions of the Order Mycoplasmatales, Family Mycoplasmataceae. Int. J. Syst. Evol. Microbiol. 43, 378–385 (1993).

50. Maier, T. et al. Quantification of mRNA and protein and integration with protein turnover in a bacterium. Mol Sys Bio 7:511 (2011).

51. Klement, M. L. R., Öjemyr, L., Tagscherer, K. E., Widmalm, G. & Wieslander, Å. A processive lipid glycosyltransferase in the small human pathogen Mycoplasma pneumoniae: Involvement in host immune response. Mol. Microbiol. 65, 1444–1457 (2007).

52. Pollack, J. D., Somerson, N. L. & Senterfit, L. B. Chemical Composition and Serology of Mycoplasma pneumoniae Lipids. J. Infect. Dis. 127, S32–S35 (1973).

53. Stoll, L. L. & Spector, A. A. Changes in serum influence the fatty acid composition of established cell lines. In vitro 20, 732–738 (1984).

54. Rottem, S. Membrane lipids of mycoplasmas. Biochim. Biophys. Acta - Rev. Biomembr. 604, 65–90 (1980).

55. Rottem, S., Adar, L., Gross, Z. V. I., Eman, N. E. & Davis, P. J. Incorporation and Modification of Exogenous Phosphatidylcholines by Mycoplasmas. J. Bacteriol. 167, 299–304 (1986).

56. Lonnfors, M., Doux, J. P. F., Killian, J. A., Nyholm, T. K. M., Slotte, J. P. & Lo, M. Sterols Have Higher Affinity for Sphingomyelin than for Phosphatidylcholine Bilayers even at Equal Acyl-Chain Order. Biophys J. 100, 2633–2641 (2011).

57. Worliczek, H. L., Kämpfer, P., Rosengarten, R., Tindall, B. J. & Busse, H.-J. Polar lipid and fatty acid profiles – Re-vitalizing old approaches as a modern tool for the classification of mycoplasmas? Syst. Appl. Microbiol. 30, 355–370 (2007).

58. Kornspan, J. D. & Rottem, S. The Phospholipid Profile of Mycoplasmas. J. Lipids 2012, 1–8 (2012).

59. Rottem, S.& Markowitz, O. Membrane Lipids of Mycoplasma gallisepticum : A Disaturated Phosphatidylcholine and a Phosphatidylglycerol with an Unusual Positional Distribution of Fatty Acids. Biochemistry. 2930–2935 (1979).

60. Low, I. E. & Eaton, M. D. Replication of Mycoplasma pneumoniae in Broth Culture. J. Bacteriol. 89, 725 LP–728 (1965).

61. Freundt, E. A. Culture media for classic mycoplasmas. Methods in Mycoplasmology I, 127–135 (1983)

62. Segovia, J. A. et al. NLRP3 Is a Critical Regulator of Inflammation and Innate Immune Cell Response during Mycoplasma pneumoniae Infection. Infect. Immun. 86, (2018).

63. Kenny, G. E., Kaiser, G. G., Cooney, M. K. & Foy, H. M. Diagnosis of Mycoplasma pneumoniae pneumonia: sensitivities and specificities of serology with lipid antigen and isolation of the organism on soy peptone medium for identification of infections. J. Clin. Microbiol. 28, 2087–2093 (1990).

64. Berglund, A. H., Nilsson, R. & Liljenberg, C. Permeability of large unilamellar digalactosyldiacylglycerol vesicles for protons and glucose – influence of α -tocopherol, -carotene, zeaxanthin and cholesterol. Plant Physiol. Biochem. 37, 179–186 (1999).

65. Osterberg, F. et al. Lipid extracts from membranes of Acholeplasma laidlawii A grown with different fatty acids have a nearly constant spontaneous curvature. Biochimica et Biophysica Acta 1257, 18–24 (1995).

66. Finkelstein, A. Water and Nonelectrolyte Permeability of Lipid Bilayer Membranes. J Gen Physiol 68, 127–135 (1976).

67. Razin, S., Kutner, S., Efrati, H. & Rottem, S. Phospholipid and cholesterol uptake by mycoplasma cells and membranes. Biochim. Biophys. Acta - Biomembr. 598, 628–640 (1980).

68. Cluss, R. G., Johnson, J. K. & Somerson, N. L. Liposomes replace serum for cultivation of fermenting mycoplasmas. Appl. Environ. Microbiol. 46(2), 370–374 (1983).

69. Schummer, U. & Gerhardt, U. W. The proton gradient across Mycoplasma membranes. Current Microbiol 5, 371–374 (1981).

70. Bergman, C., Kashiwaya, Y. & Veech, R. L. The Effect of pH and Free Mg 2 + on ATP Linked Enzymes and the Calculation of Gibbs Free Energy of ATP Hydrolysis. J. Phys. Chem. 114, 16137–16146 (2010).

