## Supplementary material for "Model-driven design allows growth of *Mycoplasma pneumoniae* on serum-free media": Supplementary File S1 - script to simulate_growth.py

```
# -*- coding: utf-8 -*-
"""
Created on Thu Jan 11 11:24:43 2018

@author: eryg9
"""

##################### IMPORT LIBRARIES ############################

import cobra
import pandas
import libsbml
from cobra import Model, Reaction, Metabolite
from cobra import flux_analysis

##################### READ MODEL ############################

model = cobra.io.read_sbml_model("iEG158_mpn.xml")

##################### SUPPLEMENT SERUM-FREE MEDIUM AND EXPORT WASTE ############################

amino_acids=['EX_thr__L_e','EX_leu__L_e','EX_cys__L_e','EX_glu__L_e','EX_asp__L_e',
             'EX_lys__L_e','EX_asn__L_e','EX_ala__L_e','EX_ile__L_e','EX_gly_e',
             'EX_trp__L_e','EX_met__L_e','EX_ser__L_e','EX_his__L_e','EX_orn__L_e',
             'EX_val__L_e','EX_gln__L_e','EX_tyr__L_e','EX_phe__L_e','EX_pro__L_e','EX_arg__L_e']

essential_components=['EX_o2_e','EX_h_e','EX_h2o_e','EX_pi_e']

essential_nutrients=['EX_chol_e','EX_pydx_e','EX_pap_e','EX_fol_e','EX_thm_e','EX_ribflv_e','EX_ptth_e','EX_nac_e']

lipids=['EX_sphng160_e','EX_sphng161_e','EX_sphng180_e','EX_sphng181_e','EX_sphng182_e','EX_sphng200_e','EX_sphng240_e',
        'EX_hdca_e','EX_hdcea_e','EX_ocdca_e','EX_ocdcea_e','EX_lneldc_e','EX_lnlncg_e',
        'EX_pc160_e','EX_pc161_e','EX_pc180_e','EX_pc181_e','EX_pc182_e','EX_pc183_e',
        'EX_chsterol_e']

nucleotides=['EX_ade_e','EX_cytd_e','EX_gua_e','EX_ura_e','EX_thym_e','EX_dcyt_e']

export_waste=['EX_lac__L_e','EX_h2o2_e','EX_ac_e','EX_nh3_e','EX_co2_e']

products=['EX_BM_e']

carbon_source=['EX_glc__D_e','EX_glyc_e']

Medium_n_Products = amino_acids + essential_components + lipids + essential_nutrients + nucleotides + export_waste + products + carbon_source 

for component in Medium_n_Products:
    model.reactions.get_by_id(component).lower_bound = -1000
    model.reactions.get_by_id(component).upper_bound = 1000

##################### SET EXPERIMENTALLY-DETERMINED CONSTRAINTS ############################

model.reactions.get_by_id('ALKP').lower_bound = 0.01
model.reactions.get_by_id('IR08208').lower_bound = 0.01
model.reactions.get_by_id('IR09954').lower_bound = 0.0003489
model.reactions.get_by_id('RNase').lower_bound = 0.007741
model.reactions.get_by_id('EX_glc__D_e').lower_bound = -5.10835836764991
model.reactions.get_by_id('EX_glyc3p_e').lower_bound = -0.127708959191248
model.reactions.get_by_id('EX_glyc_e').lower_bound = -0.127708959191248
model.reactions.get_by_id('EX_rib__D_e').lower_bound = -0.127708959191248
model.reactions.get_by_id('EX_ac_e').upper_bound = 7.4436
model.reactions.get_by_id('EX_nh3_e').lower_bound = -10
model.reactions.EX_arg__L_e.bounds = (-.25,1000)
model.reactions.EX_ade_e.bounds=(-.02, 1000)
model.reactions.EX_ura_e.bounds=(-.02,1000)
model.reactions.EX_cytd_e.bounds=(-.02,1000)
model.reactions.EX_dcyt_e.bounds=(-.02,1000)
model.reactions.EX_pap_e.bounds=(0,1000)
model.reactions.EX_gua_e.bounds=(-.02,1000)
model.reactions.EX_thym_e.bounds=(-.02,1000)
model.reactions.ALKP.bounds=(-0.01,1000)
model.reactions.G3PO.bounds = (-1000 , .1)
model.reactions.ATPsyn.bounds = (-1000 , 1000)
model.reactions.GLUN.bounds = (-0.1, 0.1 )

##################### MAINTENANCE REACTION ############################

total_maintenance = 10.46
prot_leak_fraction = .66

model.reactions.protonLeak.lower_bound = total_maintenance * prot_leak_fraction * 4
model.reactions.IR08984.lower_bound = total_maintenance * (1-prot_leak_fraction)

##################### SOLVE THE MODEL ############################

model.solver='gurobi'
growth_rate=model.optimize()
print(growth_rate)
print(growth_rate.f)

##################### SIMULATE HYPOXIA ############################

#model.reactions.EX_o2_e.bounds=(-6,0)


   
```
