## Supplementary figures and images for "Model-driven design allows growth of *Mycoplasma pneumoniae* on serum-free media"

### Supplementary Figure S1.jpg

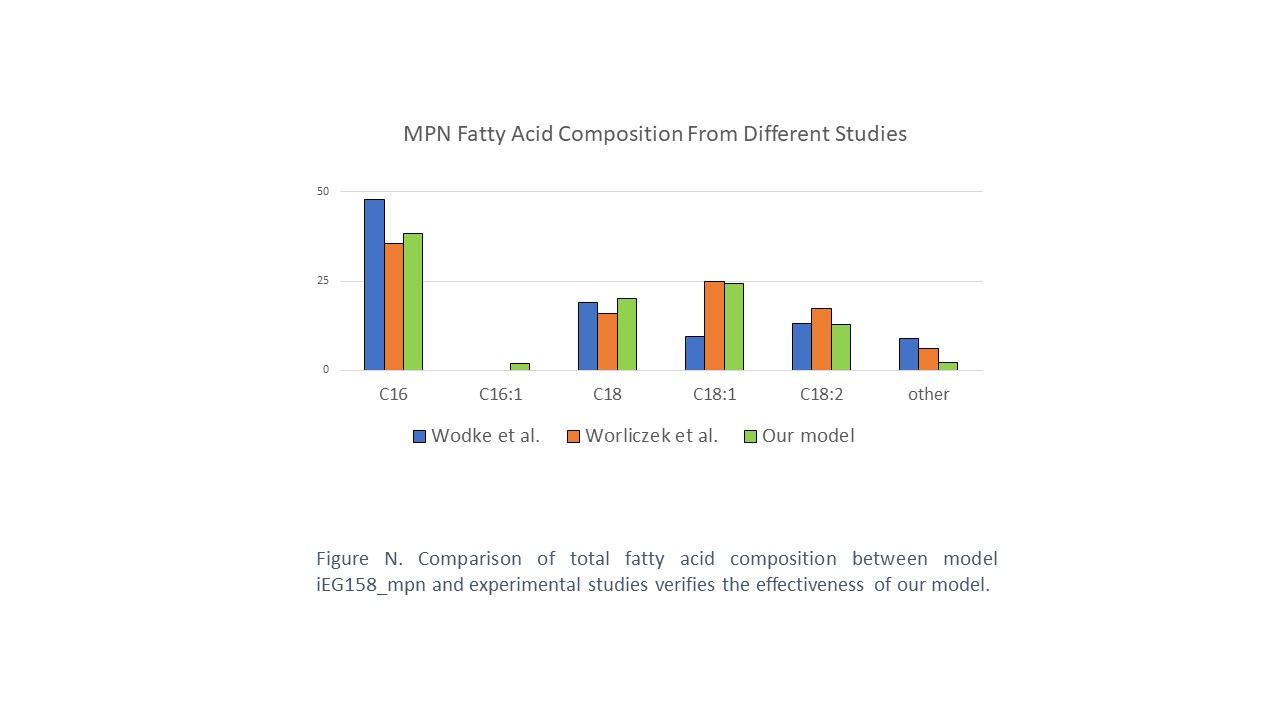
